# A Method for Cost-Effective and Rapid Characterization of Genetic Parts

**DOI:** 10.1101/2021.04.30.440836

**Authors:** John B. McManus, Casey B. Bernhards, Caitlin E. Sharpes, David C. Garcia, Stephanie D. Cole, Richard M. Murray, Peter A. Emanuel, Matthew W. Lux

## Abstract

Characterizing and cataloging genetic parts are critical to the design of useful genetic circuits. Having well-characterized parts allows for the fine-tuning of genetic circuits, such that their function results in predictable outcomes. With the growth of synthetic biology as a field, there has been an explosion of genetic circuits that have been implemented in microbes to execute functions pertaining to sensing, metabolic alteration, and cellular computing. Here, we show a cost-effective and rapid method for characterizing genetic parts. Our method utilizes cell-free lysate, prepared in-house, as a medium to evaluate parts via the expression of a reporter protein. Template DNA is prepared by PCR-amplification using inexpensive primers to add variant parts to the reporter gene, and the template is added to the reaction as linear DNA without cloning. Parts that can be added in this way include promoters, operators, ribosome binding sites, insulators, and terminators. This approach, combined with the incorporation of an acoustic liquid handler and 384-well plates, allows the user to carry out high-throughput evaluations of genetic parts in a single day. By comparison, cell-based screening approaches require time-consuming cloning and have longer testing times due to overnight culture and culture density normalization steps. Further, working in cell-free lysate allows the user to exact tighter control over the expression conditions through the addition of exogenous components, or by titrating DNA concentrations rather than relying on limited plasmid copy numbers. Because this method retains a cell-like environment, the function of the genetic part will typically mimic its function in whole cells.

**SUMMARY:** Well-characterized genetic parts are necessary for the design of novel genetic circuits. Here we describe a cost-effective, high-throughput method for rapidly characterizing genetic parts. Our method reduces cost and time by combining cell-free lysates, linear DNA to avoid cloning, and acoustic liquid handling to increase throughput and reduce reaction volumes.

## INTRODUCTION

A core effort of synthetic biology is the development of genetic tool kits containing well-characterized parts, which can be used to construct genetic circuits^1^ that carry out useful functions when deployed in microbes or cell-free lysate. Areas in which such genetic circuits have found purchase are sensing^2–5^, human performance^6,7^, biofuels^8,9^, materials production^10,11^, and cellular computing^12^. Registries of standardized genetic parts have been established^13^ to catalog new and existing parts into categories like promoters, operators, coding sequences, and terminators, to name just a few. Efforts such as the iGEM competition^14^ have been instrumental in characterizing and cataloging these genetic parts. Methods, such as UNS^15^ and 3G^16^, have been developed to facilitate the rapid assembly of these parts into useful genetic circuits. Software, such as Cello^17^, have even been developed to automate the composition of well-characterized parts into circuits that achieve a desired function. However, the assembly of useful genetic circuits with predictable function rests on the presumption that the genetic tool kits contain well-characterized genetic parts. Due to the necessity of these tool kits toward the advancement of synthetic biology, numerous undertakings to better catalog circuits and parts with appropriate characterization data have been described^18–22^.

One category of components useful to the implementation of genetic circuits are orthogonal parts, such as T7 RNA polymerase (T7 RNAP) and its cognate T7 promoter. The genetic systems of *Escherichia coli* are well-developed and many genetic circuits have been deployed and characterized in this organism. T7 RNAP is particularly well-suited as an orthogonal actuator for genetic circuits in *E. coli*, owing to its ability to partially insulate circuit function from the host metabolism^23^. However, the major drawback of T7 RNAP is the lack of direct regulatory mechanisms. Until recently, regulated T7 promoters were limited to a small set that had been engineered by insertion of bacterial operator sequences^24,25^. In order to fill this gap, we wanted to develop a method for the rapid characterization of libraries of regulated T7 promoters. We originally developed the method, presented here, for rapidly characterizing spatial combinations of the T7 promoter and bacterial operator sequences along with their cognate repressor proteins. We validated our methodology using the tetracycline regulatory system (*tet*), by investigating the effects of proximity of the tetracycline operator (*tetO*) sequence to the T7 promoter on the regulation of T7 RNAP-driven expression. Our results revealed important insights into the kinetics of T7-driven expression and into the future design of engineered T7-based transcription factors^26^. We believe that this methodology can be applied to the characterization of other types of genetic parts as well. In the meantime, others have expanded the range of such regulated T7 promoters considerably^27^.

Here, we present methodology that uses PCR to amplify template DNA via primers that add variant genetic parts to a reporter gene. These genetic parts are evaluated using cell-free lysate, prepared in-house, as a cost-effective medium to measure the expression of the reporter protein by cell-free protein synthesis (CFPS). Several studies have demonstrated the utility of CFPS for prototyping genetic components^28–32^. Note that while we prepare our cell-free lysates here based on Sun et al.^33^, numerous other commercial kits and protocols^34,35^ should work similarly. We have included the use of an acoustic liquid handler and 384-well plates to increase throughput and decrease the volume of materials required. Previous work has demonstrated successful use of acoustic liquid handling at significantly lower volumes^36,37^ with variability comparable to manual pipetting at larger volumes^38^. CFPS removes the requirement of working in whole cells and reduces the amount of time for screening many genetic parts to a single day, while still maintaining a cell-like environment. A second advantage of CFPS is that the user can exact tighter control over the expression conditions through the addition of exogenous components via the acoustic liquid handler. The generation of variants by changing PCR primers and then using linear DNA in the CFPS reactions dramatically cuts the cost and time of variant preparation compared to cloning or synthesis. Though we had initially developed this method for the characterization of engineered T7 promoters regulated by transcription factors^26^, other parts that can be changed by PCR-amplification include promoters, operators, RBS sequences, insulators, and terminators. We hope that through this methodology, the synthetic biology community can grow the number of characterized parts for the assembly of predictable genetic circuits with useful function.

## PROTOCOL

1. Preparation of Cell Extract

1.1. Preparation of Media

1.1.1. For 2xYT media: Add 16 g of tryptone, 10 g yeast extract, 5 g NaCl to 900 mL of deionized water. Adjust pH to 7.0 with 5 M NaOH and adjust volume of solution to 1 L using deionized H2O. Alternatively, purchase 2xYT media.
1.1.2. For S30B buffer: Prepare a solution of 14 mM Mg-glutamate, 60 mM K-glutamate, and 5 mM Tris (pH 8.2) in 2 L of deionized water. Use 2 M Tris to get pH to 8.2. Store at 4 °C. Complete solution by adding DTT to 1 mM final concentration just before use.
1.2. Preparation of Cells

1.2.1. Streak *E. coli* K12 Rosetta cells onto a plate of LB agar and incubate at 37°C for 10 to 14 h.
1.2.2. From a single colony, inoculate one 10 mL culture tube containing 3 mL of 2xYT medium with *E. coli* K12 Rosetta cells. Incubate this tube at 37°C, shaking at 250 rpm, for 8 h.
1.2.3. From the 3 mL culture, inoculate 50 μL into a 500 mL flask containing 50 mL 2xYT medium. Incubate this flask at 37°C, shaking at 250 rpm, for 8 h.
1.2.4. From the 50 mL culture, inoculate 7.5 mL into four 4 L baffled flasks containing 0.75 L 2xYT medium. Incubate these flasks at 37°C, shaking at 220 rpm, until they have reached an optical density at 600 nm of 2 to 4, approximately 3-4 h.
1.2.5. Harvest the cells from each flask by centrifugation, in 1 L containers, at 5000 × g for 12 min. Discard the supernatant.
1.2.6. Wash each cell pellet with 150 mL of ice cold S30B buffer by completely resuspending them, then collect the cells again by centrifugation at 5,000 × g for 12 min. Discard the supernatant.
1.2.7. Wash each cell pellet again in 40 mL of ice cold S30B buffer by completely resuspending them. Transfer the cells to a 50 mL conical tube and collect the cells again by centrifugation at 775 × g for 8 min. Discard the supernatant.
1.2.8. Weigh the cell pellets. Flash freeze the cell pellets in liquid nitrogen. Store the cell pellets at −80°C.
1.3. Cell Lysis

1.3.1. Thaw cell pellets on ice.
1.3.2. Resuspend each cell pellet in 1.4 mL of S30B buffer per 1 g of cell pellet.
1.3.3. Lyse the cells by French pressure cell at 640 psi at 4°C. Collect the lysate on ice and add 3 μL of 1 M DTT per 1 mL of lysate immediately after lysis. NOTE: It is best to tap the French press release valve with a small metal rod in order to maintain even pressure and avoid sudden drops in pressure.
1.3.4. Clear the lysate by centrifugation at 30,000 × g for 30 min at 4°C and discard the pellet.
1.3.5. Centrifuge the supernatant a second time at 30,000 × g for 30 min at 4°C and discard the pellet.
1.3.6. Incubate the supernatant in a 37°C water bath for 1 h.
1.3.7. Clear the supernatant by centrifugation at 15,000 × g for 15 min at 4°C and discard the pellet.
1.3.8. Centrifuge the supernatant a second time at 15,000 × g for 15 min at 4°C and discard the pellet.
1.3.9. Distribute the supernatant in 100 μL aliquots into 1.5 mL microcentrifuge tubes and flash freeze them in liquid nitrogen. Store the supernatant at −80°C.
2. Linear Template Preparation

2.1. Primer Design

2.1.1. For the forward primers, choose a minimum of 20 bp on the 3⍰ end of the primer to match the 5⍰ end of the coding strand of the reporter gene. Design the remainder of the 5⍰ end of the primer to add the genetic parts of interest to the reporter gene via PCR-amplification (Fig. 1A and Fig. 2).
2.1.2. For the reverse primer, choose a sequence on the 5⍰ end of the non-coding strand of the reporter gene, directly downstream from the terminator. Alternatively, for parts at the end of the gene, such as C-terminal tags, design primers to introduce the part of interest. Note that inclusion of a terminator may not be required for expression from linear DNA in cell-free systems. Be sure that the reverse primer’s annealing temperature is within 5°C of the annealing temperature of the entire forward primer. NOTE: DNA providers typically synthesize primers with a standard length of up to ~60 bases. While longer primers can be synthesized and implemented, costs often increase dramatically. Alternatively, multiple overlapping primers can be designed to the 5⍰ or 3⍰ ends of the gene, and longer sequences or multiple parts can be added to the reporter through multiple rounds of PCR. NOTE: The template containing the reporter gene to be amplified need not be a plasmid. Genomic DNA or linear blocks ordered from commercial vendors may also be suitable as template DNA.
2.2. Linear Template Amplification

2.2.1. Determine the number of PCR reactions to perform and calculate the amount of each component required using Table 1.
2.2.2. Prepare the master mix and store it on ice. Aliquot 40 μL of the master mix into the determined number of PCR tubes and add 10 μL of Forward Primer (5 μM) to each appropriately-labeled, corresponding PCR tube.
2.2.3. Place PCR tubes into the thermocycler and run the following PCR program:

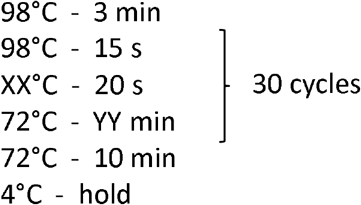 Where XX represents the temperature for the lower annealing temperature primer. YY represents the extension time calculated for the length of the amplicon (30 s – 1 min / kb for Q5 polymerase). These conditions may need to be optimized for different primers and/or templates (see 2.3 and Discussion).
2.2.4. Add 1 μL of NdeI restriction enzyme to digest the original template. Incubate the reaction at 37°C for 1 h. This step is not necessary if a synthetic block was used as the original template.
2.2.5. Analyze 5 μL of each PCR product by gel electrophoresis. Separate the product using a 1% agarose gel at 180 V for 20 min.
2.3. Purify the linear template according to the QIAquick PCR Purification Kit, or by your preferred PCR cleanup method. If multiple bands were present by gel electrophoresis analysis, see the Discussion section for troubleshooting OR purify the correct molecular weight bands by QIAquick Gel Extraction Kit, or by your preferred extraction method.
2.4. Quantify each DNA template using a NanoDrop spectrophotometer, or your preferred spectrophotometric instrument. Be sure to assess DNA template quality by assuring that the 260 nm / 280 nm ratio is approximately 1.8. Further, a portion of the samples can be separated using a 1% agarose gel at 180 V for 20 min, a second time, to check that any unwanted bands were removed during template purification. DNA samples may be stored at −20 °C.
3. Purified Protein Preparation

3.1. Protein Expression

3.1.1. For each protein to be expressed, assemble the *E. coli*-codon optimized gene into a pET-22b expression vector and transform the expression plasmid into BL21(DE3) Rosetta expression cells.
3.1.2. For each protein, inoculate, from a single colony, a 10 mL culture tube containing 3 mL of LB medium. Incubate these tubes at 37°C, shaking at 250 rpm, overnight.
3.1.3. Inoculate a 2 L flask containing 750 mL of LB medium with 1 mL of overnight culture. Incubate these flasks at 37°C, shaking at 250 rpm, until they reach an OD_600_ of 0.6–1.0.
3.1.4. Induce protein expression by adding 0.75 mL of 1 M IPTG in water to each flask, and continue to incubate these flasks at 37°C, shaking at 250 rpm, for 4 h.
3.1.5. Harvest the cells from each flask, using a 1 L centrifuge bottle, by centrifugation at 5000 × g for 12 min. Discard the supernatant.
3.1.6. Transfer the pellets to a 50 mL conical tube and weight each pellet. Expect 2-5 g per 0.75 mL. Flash-freeze the cells in liquid nitrogen and store them at −80 °C or proceed to 4.2.1.
3.2. Protein Purification by Nickel Affinity Column Chromatography

3.2.1. Thaw the cell pellet in room temperature water and resuspend, homogenously, in lysis buffer, using 5 mL lysis buffer per 1 g cell pellet.
3.2.2. Lyse the cells by sonication. Separate the cell homogenate so that there is no more than 30 mL per 50 mL conical tube and place each tube on ice. Lyse the cells using a QSonica Ultrasonic Processor sonicator with a 0.16 cm diameter probe. Lyse the cells in 15 s rounds with 30 s breaks, 10 times. NOTE: Avoid foaming, as this denatures protein. Formation of foam can be avoided by keeping the tip at least 2/3 submerged in the lysate while it is operational.
3.2.3. Clear the lysate by centrifugation at 15,000 × g for 30 min at 4 °C.
3.2.4. Incubate each 5 mL of supernatant with 1 mL of Ni-NTA resin at 4 °C, on a Thermo tube rotator (or equivalent) at 10 rpm for 1 h. Separate the cell lysate / Ni-NTA slurry so that there is no more than 36 mL per 50 mL conical tube.
3.2.5. Load the resin into a 5 cm diameter column. Wash the resin with 10 resin bed volumes of wash buffer.
3.2.6. Collect the protein with three resin bed volumes of elution buffer and concentrate the volume to 1.5 mL using a centrifugal concentrator with the appropriate molecular weight cut-off membrane for each protein.
3.2.7. Dialyze the protein against 2 L of dialysis buffer at 4 °C for 1 h. Dialyze the protein again against 2 L of dialysis buffer overnight at 4 °C.
3.2.8. Quantify the protein using its molar extinction coefficient and absorbance at 280 nm. Analyze the protein for purity by separating it using SDS-PAGE electrophoresis. Store the protein at −80 °C.
4. Cell-Free Protein Synthesis

4.1. Preparation of CFPS Reaction Mixture

4.1.1. Prepare Supplement Mix by following the Amino Acid Solution Preparation, Energy Solution Preparation, and Buffer Preparation steps in Sun et al.^33^ These three solutions can be combined ahead of time, aliquoted, and stored at −80 °C. Final concentrations should match those described in Sun et al.^33^ in the Experimental Execution of a TX-TL Reaction section.
4.1.2. Prepare GamS (protects linear DNA from degradation; see note in next section), T7 polymerase, and repressor proteins using steps above (see 3. Purified Protein Preparation), or obtain from a commercial vendor.
4.1.3. Determine the number of CFPS reactions to be performed and calculate the amount of each component required using Table 2. NOTE: The table above is a for a standard CFPS reaction. The volume of the template, polymerase, and repressor protein can be varied or other components can be added (or subtracted) by adjusting the amount of water such that the final volume of each reaction mixture is always 10 μL. Additionally, other components can be optionally dispensed by acoustic liquid handling instead of being present in the master mix (see Troubleshooting). NOTE: While GamS protein is used here to limit degradation of linear DNA by blocking the RecBCD complex^39,40^, other approaches are available^41–43^ and some CFPS recipes based on purified components do not require any additions^44^.
4.1.4. Thaw all components on ice and prepare a master mix by mixing each component as calculated above. Mix all the components thoroughly, by pipette. Keep the master mix on ice.
4.1.5. Chill a 384-well plate on ice and distribute the master mix in 9 μL aliquots into each well using an electronic repeater pipette.
4.2. Distribution of Additional Components by Acoustic Liquid Handling

4.2.1. Calculate the amount of repressor protein (and optional other components) required for all the CFPS reactions.
4.2.2. Thaw the repressor protein on ice and distribute it into a Labcyte Echo source plate or other appropriate plate. Be sure to consider the appropriate amount of dead volume required for the type of source plate used.
4.2.3. Distribute the repressor protein in 1 μL volumes into the appropriate wells via the Echo acoustic liquid handler or similar instrument. More information on distribution troubleshooting can be found in the Discussion sections.
4.3. Standard Curves

4.3.1. Include a serial dilution of purified reporter protein (see Section 3 for protein purification)^38^] or appropriate chemical standard^45^ on the plate to enable comparison of results with other studies and other labs. In our previous publication^26^, a standard curve for sfGFP was generated using purified protein with concentrations ranging from 0 μM to 10 μM.
4.4. Running CFPS Reactions

4.4.1. Pre-warm the plate reader (Biotek H10 used in our previous publication^26^) to 37°C. Ensure that the settings match those for the reporter protein being measured (Ex: GFP, fluorescence, 480 nm ex / 528 nm em). No shaking steps are required. It may be helpful to run a test reaction first to set the appropriate gain or sensitivity setting that will capture the change in fluorescence without signal overflow. NOTE: Read intervals of 10 minutes are sufficient to achieve good resolution on sfGFP expression curves using the Biotek H10 plate reader. However, this may change depending on the reporter protein and particular CFPS recipe.
4.4.2. Seal the 384-well plate with an impermeable plastic sealable lid to prevent evaporation. The instrument should be set to a 1 °C vertical temperature gradient. This ensures that the condensation does not form on the seal. Place the 384-well plate on the plate holder and begin reading.
5. Data Interpretation

5.1. At the completion of the run, export the data as a .csv file.
5.2. Data should be converted to standardized units if applicable (see Section 4.3).
5.3. For characterization of repressible promoters, a dose response curve of reporter expression against a titration of repressor concentration is informative. To interpret these data, plot the maximum output values from each reaction against the repressor protein concentration, then fit each dose response profile to a four-parameter logistic curve using GraphPad Prism or the software of your choice. NOTE: The maximum output values used here are derived from the sigmoidal fit of each protein expression curve.
5.4. Calculate the maximum repression values for each data set by subtracting the top and bottom values, determined by the four-parameter logistic curve fit. The maximum repression values and EC_50_ values, determined by the four-parameter logistic fit, can be used to identify changes in the operator-promoter relationship.

**Figure 1.**
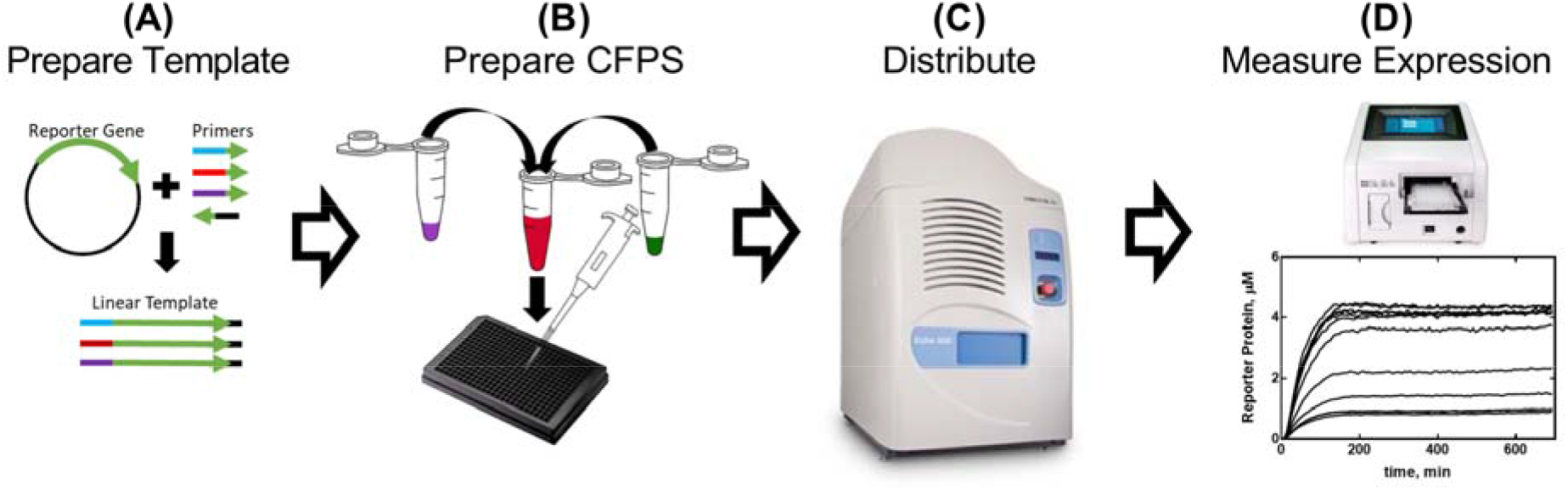
Single-day Workflow for the Evaluation of Promoter Parts in Cell-Free Extract. **(A)** A reporter gene is PCR-amplified using primers containing genetic parts to be evaluated in order to prepare template for CFPS reactions (2–5 h). **(B)** The cell-free reaction mix is prepared as detailed in the Protocol and distributed into a 384 well plate (30 min). The template can be added into the cell-free master mix or as part of the **(C)** distribution of additional components, which can include repressor proteins, effector molecules, and any other conditional effectors (10 min). **(D)** Reporter protein expression from each reaction is measured in a plate reader (2 – 16 h, depending on CFPS recipe and construct).

**Figure 2.**
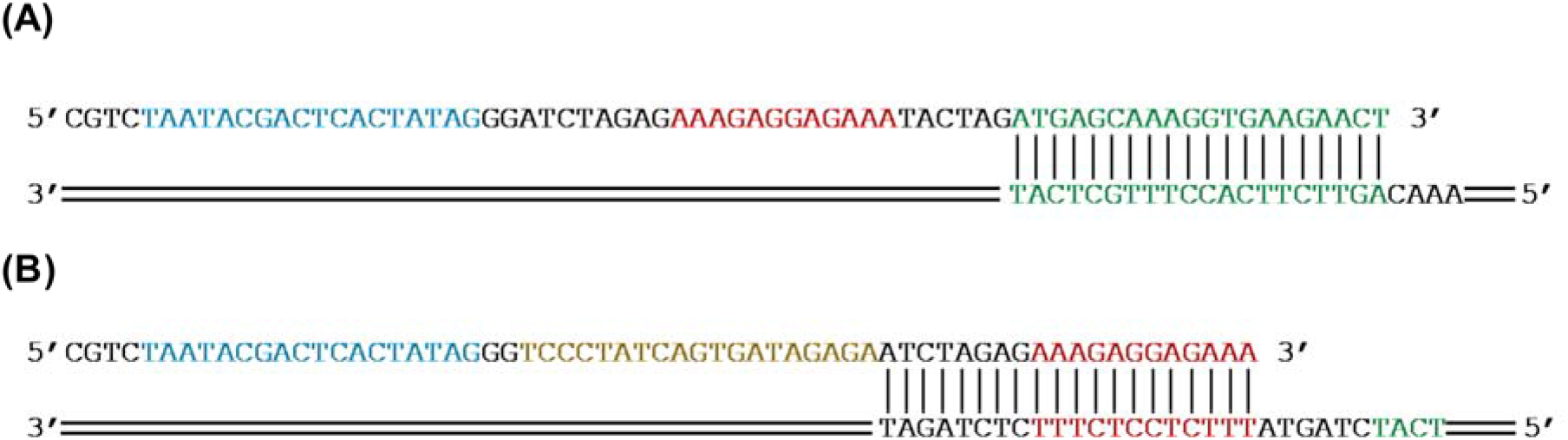
Primer Design for Adding Genetic Parts to a Reporter Gene by PCR-Amplification. Design the 3⍰ end of the primer (top) such that it will hybridize to at least 20 bp of the 3⍰ end of the non-coding strand of the reporter gene (bottom). **(A)** The sfGFP reporter gene (green) will be amplified to add an RBS (red) and a T7 promoter (blue) by PCR. **(B)** The sfGFP (green) AND an RBS (red) will be amplified to add a tetO sequence (gold) and a T7 promoter (blue) by PCR.

**Table 1.** Reagents for PCR reactions.

| Component Name | Volume for 1 reaction (μL) | Volume for 110% of X number of reactions (μL) |
| --- | --- | --- |
| Q5 PCR Premix | 25 |  |
| Water | 4 |  |
| Reverse Primer (5 μM) | 10 | |
| Template (1–3 ng/μL) | 1 | |
| **Master Mix Total:** | 40 |  |
| Forward Primer (5 μM) | 10 | |

**Table 2.** Reagents for CFPS reactions.

| Component Name | Volume for 1 reaction (μL) | Volume for 110% of X number of reactions (μL) |
| --- | --- | --- |
| Cell Extract | 4.2 |  |
| Supplement Mix | 3.3 |  |
| GamS Protein (207 μM) | 0.15 | |
| Template DNA (20 nM) | 1 |  |
| T7 Polymerase (13 mg/mL) | 0.12 |  |
| Water | 0.73 (this number may vary) |  |
| **Master Mix Total:** | 9 |  |
| Repressor Protein: | 1 |  |

## REPRESENTATIVE RESULTS

### TetO Represses T7 RNAP within the First 13 Bases Downstream from the T7 Promoter

To demonstrate the utility of our methods, we present results that describe the effects of proximity of the *tetO* sequence to the T7 promoter on the regulation of T7 RNAP-driven expression. The full results and their implications can be found in the work of McManus et al.^26^. Fifteen linear templates, varying only in the distance of the T7 promoter relative to the *tetO* sequence, were prepared by PCR-amplifying the sfGFP reporter as described in Linear Template Preparation (see Section 2). Amplicons were analyzed by gel electrophoresis and added to CFPS reactions, then distributed into a 384-well plate. sfGFP expression was measured from each template with a titration of 12 different concentrations of the TetR protein, in triplicate, using an Echo acoustic liquid handler. At 36 CFPS reactions per template and 15 templates, a total of 540 reactions for the entire set of T7- *tetO* combinations were performed. The entire evaluation was carried out on two plates in two Biotek H10 plate readers.

For data analysis, the sfGFP expression curves were first converted to units of μM sfGFP via standard curve, then fit to a sigmoidal regression. The maximum expression values as determined by the sigmoid fit were plotted against TetR concentration, and these dose-response profiles were fit to a four-parameter logistic to yield EC_50_ values and maximum repression values as described in the Protocol. We found that expression values varied from template to template. This may be in part due to impurities carried over during template purification. However, as shown in our previous publication^26^, we were able to normalize these expression values and derive meaningful conclusions by comparing these normalized values. Analysis of these values shows that the T7 RNAP downregulates T7-driven expression equally up through 13 bp downstream from the start of the T7 transcript (Fig. 3). This has implications for the future design of regulatable T7-driven gene circuits.

**Figure 3.**
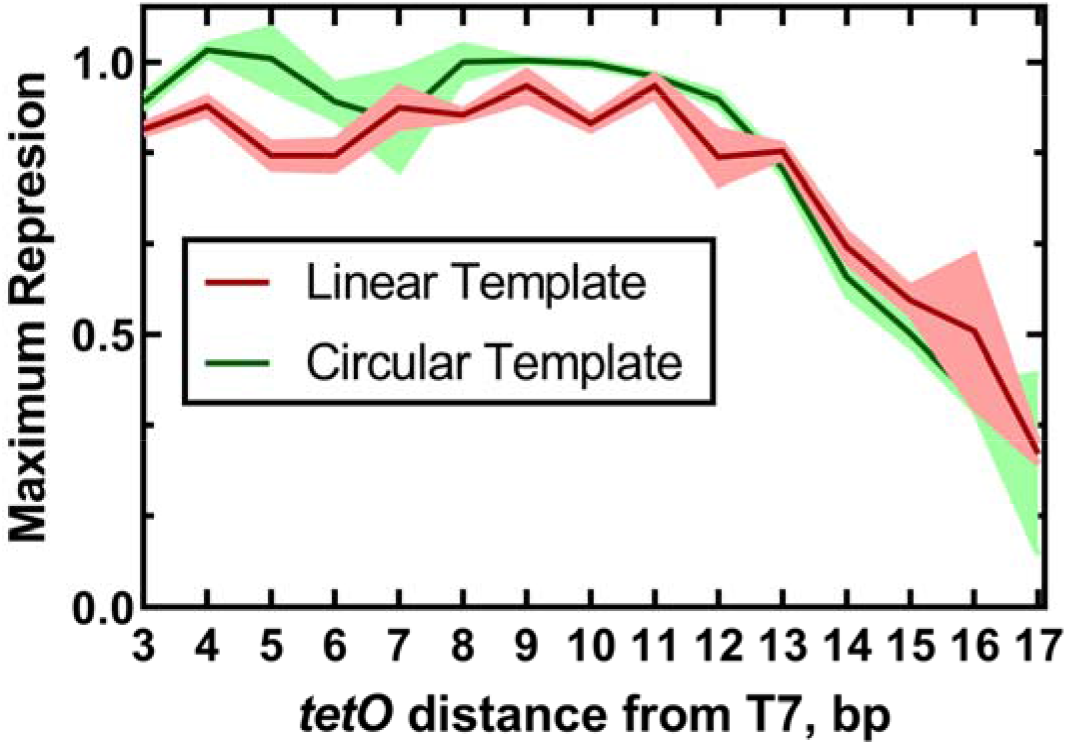
The Effect of tetO Position on the Regulation of T7-Driven Expression. Maximum, normalized repression values were plotted against tetO position to reveal the effects of tetO position on the regulation of T7-driven expression. The traces represent the mean and standard deviations for three replicates. This figure has been modified from McManus et al.^26^ under a Creative Commons CC-BY license.

We also performed the same experiment using circular plasmid DNA as template, rather than PCR-amplified template. The purpose of this experiment was to determine the differences, if any, between circular and linear template. Our results, detailed in our previous manuscript^26^, indicated that while the pattern of regulation, presented in the previous section, remains unchanged (Fig. 3), the EC_50_ values show a significant difference. We hypothesized that non-specific binding of TetR to the vector DNA could explain the observed difference. Experimental results showed that addition of linear vector DNA to reactions with linear template DNA reduced the difference to non-statistical significance, though did not rule out contributions from other factors, such as differences in periodicity of the DNA helix for linear vs. circular formats, which, in turn, could affect TetR binding. Depending upon the application, the use of linear template may require additional validation.

## DISCUSSION

The protocols described here provide a cost-effective and rapid means to screen genetic parts via the expression of a reporter protein by CFPS. Well-characterized genetic parts are crucial to the design of predictable genetic circuits with useful function. This methodology increases throughput and decreases the time needed to screen new genetic parts by removing the requirement to work in living cells, while retaining functionality that mirrors the cellular environment by retaining the metabolic process of protein expression in the cell lysate. Our protocol can performed in one day after receipt of primers, compared to at least three for traditional cloning (one day each for construct assembly and transformation, sequence verification of clones, and culturing of cells for assessment). We further estimate the cost per construct to be roughly one third compared to traditional cloning (Supplementary Table 1). Commercial synthesis services take a minimum of 5 business days, though may have similar costs to our methods if linear fragments are screened directly in CFPS; we have not verified this approach.

### Customizing the Methodology

We originally developed this methodology to investigate the effects of operator proximity on the T7 promoter^26^. We have attempted to present the protocols here in a more generic format, such that they can be applied to promoters, operators, ribosome binding sequences, insulators, and terminators. These genetic parts can be added to the 5⍰ or 3⍰ end of the reporter gene by PCR using primers for each design, obviating the need for synthesis or cloning of each variant to test. The resulting PCR products serve as template DNA for evaluation via the expression of a reporter protein. In our work, the affinity purification protocol provided here was used for TetR and GamS. The same procedure can be used for the expression and purification of other repressors, activators, polymerases, sigma factors, and other proteins cognate to a genetic part of interest. Purification and titration of these proteins into CFPS reactions enables a more detailed characterization of a particular genetic part. The procedure given here for purifying protein may need to be altered to the desired protein being expressed. Finally, numerous alternative CFPS protocols exist and each should be amenable to this methodology. Varying the concentrations of underlying constituent components of the CFPS is also possible. The use of acoustic liquid handling enhances the ability to tests the myriad conditions by increasing throughput and decreasing materials required.

### Future Directions

Beyond characterization of individual parts, the same method can be used to screen combinations of parts that form complex circuits, such as logic circuits^17^ or oscillators^46,47^. Alternatively, the method can be used for screening and optimization of biosensors for applications in epidemiological diagnostics^48–51^ or hazard detection and quantification^3,4,52^. Similar to our TetR-responsive T7 promoters study, these detection circuits are frequently designed around the interaction between a repressor protein and its cognate effector in order to regulate the expression of genes via an operator sequence, making our method highly appropriate; however, other biosensor mechanisms, such as riboswitches, can also be screened with our approach. Further, the *E. coli* cell-free lysate can also be replaced with lysate from the organism in which a biosensor is to be deployed in order to better mimic the function of that sensor in its host organism. Several publications have described the preparation of different cell lysates for protein expression including those from *Bacillus subitilis*^53^, *Pseudomonas putida*^54^, and *Vibrio natrigens*^55^, to name a few. Their methodology can be used in place of the Preparation of Cell Extract (1) protocol, if desired.

### Troubleshooting

Finally, it is worth noting that different applications may require different levels of optimization before the collection of useful experimental data can begin. Four key areas are: (1) in the PCR-amplification of template, (2) in the distribution of components by the acoustic liquid handler, (3) protein expression and purification, and (4) CFPS performance.

Depending on the length and identity of the primers used to amplify linear template, the conditions of the PCR reaction may have to be optimized to increase the amount of template synthesized. Some problems users may encounter are: (a) weak or no amplification of the desired product, (b) primer-dimer formation, and (c) products resulting from amplification due to non-specific binding. It is helpful to check the primers, before ordering them, to ensure that they do not form primer-dimers. There are several web-based tools (ex: idtDNA) that can be accessed to analyze primer pairs for primer-dimer formation. If little or no product is produced, applying a temperature gradient at the annealing step may be useful in order to optimize this step. If the user is still unable to generate product in this way, consider redesigning the primers that hybridize with a new sequence on or near the reporter gene. Finally, the synthesis of products due to non-specific amplification can be reduced or eliminated by thermocycling methods like touchdown PCR^56^. Again, redesigning primers may also eliminate the problem of non-specific amplification.

Acoustic liquid handler dispensing should be optimized for each component being transferred and it is strongly recommended to run controls to verify proper distribution and reproducibility before collecting data. The ideal source plate type and liquid class setting will depend on the specific liquid to be dispensed and its components. It is not recommended to use Echo Qualified 384-Well Polypropylene 2.0 Plus Microplates to dispense DNA, as the amine coating may interact with the DNA. It should also be noted that the ability to dispense higher concentrations of certain components may depend on the acoustic liquid handler model. For example, the Echo 550 model is able to dispense liquids containing high concentrations of salts and DMSO, whereas these components must be at lower concentrations for successful transfer with the Echo 525. A test liquid transfer may be conducted by dispensing onto a foil plate seal to visualize successful droplet formation; however, this test provides limited information and droplets from different settings may appear identical. The use of a water-soluble dye, such as tartrazine, may be used to more accurately verify the correct volume is dispensed with a given setting or workflow (Fig. 4). Optimal Echo programming of liquid transfers can also influence the accuracy and consistency of data generated; for transfers > 1 μl from one source well to one destination well, sequential transfers of ≤ 1 μl should be programmed to reduce systematic well-to-well variability (Fig. 4). Lastly, theoretical and actual source well dead volumes can vary dramatically depending on source plate type, liquid class setting, and components of the specific liquid; using the Echo to survey the well volumes prior to running a program may help to gauge how accurately the Echo is able to measure a particular liquid.

**Figure 4.**
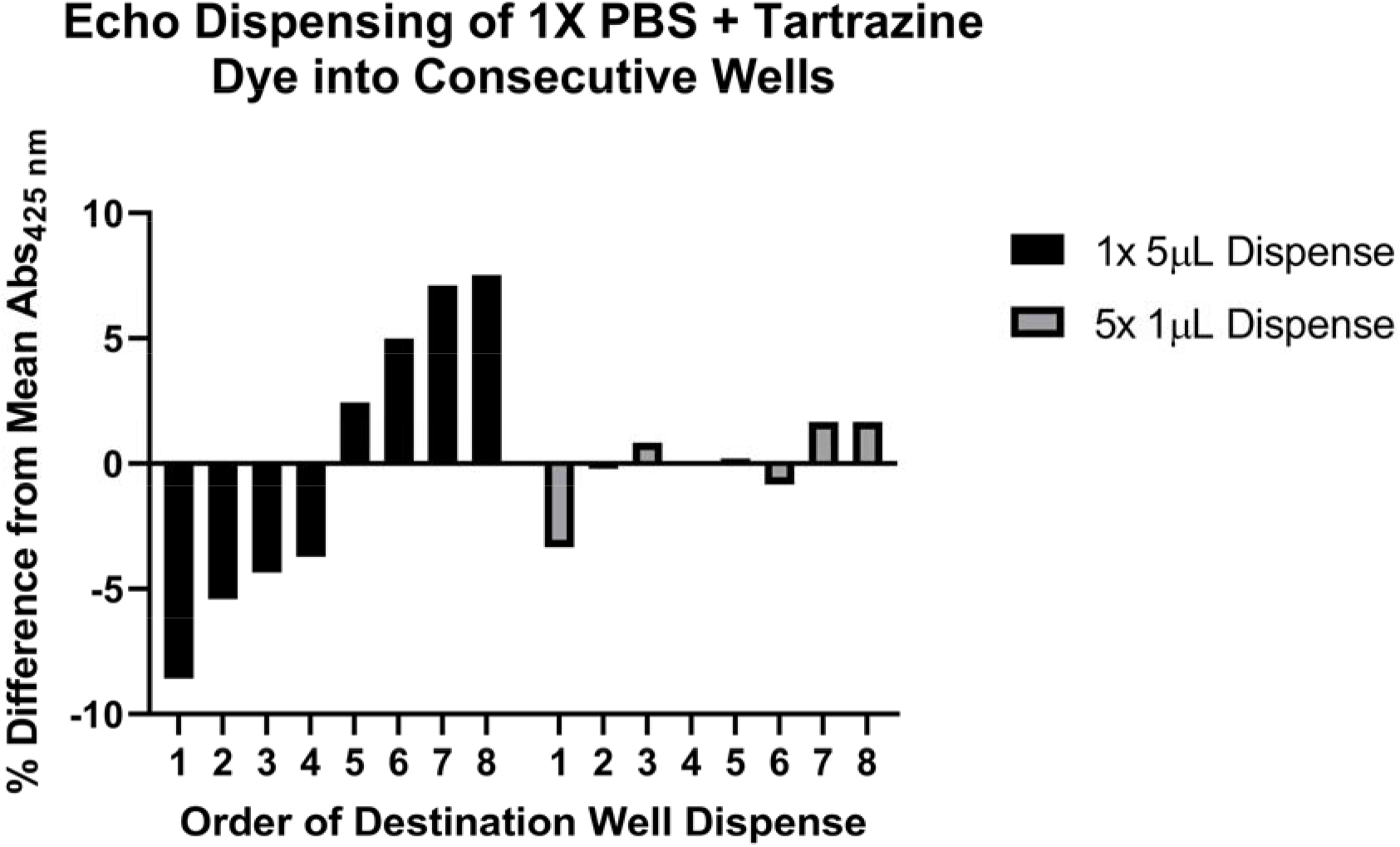
Using Tartrazine Dye to Validate Liquid Dispensing with an Acoustic Liquid Handler. A solution of 1x phosphate buffered saline (PBS), pH 7.4 containing 0.25 mM tartrazine dye was used to evaluate two methods of programming an Echo acoustic liquid handler to dispense volumes > 1 μl. In one method (results shown in black bars), 5 μl of the tartrazine solution were dispensed from a single source well into each of eight consecutive destination wells of a 384-well plate using a single programming command. In a second method (results shown in gray bars), 1 μl was dispensed from a single source well into each of eight consecutive destination wells using a single programming command, and then this step repeated a total of five times. The destination plate was sealed and centrifuged at 1,500 × g for 1 min, and the absorbance at 425 nm measured with a plate reader. Representative results of nine experiments are shown and demonstrate more consistent dispensing across the series of eight destination wells when the 5 μl transfer is divided into separate 1 μl dispenses. While the theoretical final volume in each destination well is the same for both methods (5 μl), the second method utilizes five separate programming commands resulting in the Echo resurveying the source well between each command. Based on these observations and personal communications with Labcyte engineers, it is recommended that transfers > 1 μl be broken down into multiple transfers of ≤ 1 μl to improve accuracy.

CFPS reaction performance can vary when comparing results between different users, plate readers, and laboratories^38^. For instances where such comparisons are required while prototyping genetic circuits, we recommend including internal control reactions with standard constitutive promoters (such as pT7) in each reaction plate to help normalize results across experimental set-ups. The method of DNA preparation can also contribute majorly to CFPS activity; inclusion of an ethanol precipitation step is recommended. In addition, it is useful to validate certain aspects of the reaction preparation. For instance, it has been noted that optimal magnesium and potassium glutamate concentrations can vary depending on the promoter or reporter protein being expressed^29,57^ or per batch of cell extract prepared^33^. Concentrations of these components should be optimized by screening across several concentrations of each component per genetic construct and per cell extract preparation to determine the optimal conditions for protein expression. Finally, best practices for consistent CFPS reaction performance include thorough mixing, careful pipetting, and consistency in the preparation of each reagent component.

Though this section does touch on some more common issues that may arise during screening, it is by no means an exhaustive list. Once the proper controls have been run to ensure the collection of usable and reproducible data, the method developed here is an ideal way to rapidly characterize genetic parts with a high degree of resolution.

## ACKNOWLEDGMENTS

This work was made possible by the Office of the Secretary of Defense’s Applied Research for the Advancement of Science and Technology Priorities program. We thank Scott Walper (Naval Research Laboratory) for providing the stock of sfGFP used, and Zachary Sun and Abel Chiao (Tierra Biosciences) for fruitful discussions related to prototyping with cell-free systems and related troubleshooting of acoustic liquid handling.

